# Comprehensive transcriptional profiling of the gastrointestinal tract of ruminants from birth to adulthood reveals strong developmental stage specific gene expression

**DOI:** 10.1101/364752

**Authors:** S. J. Bush, M. E. B. McCulloch, C. Muriuki, M. Salavati, G. M. Davis, I. L. Farquhar, Z. M. Lisowski, A. L. Archibald, D. A. Hume, E. L. Clark

## Abstract

One of the most significant physiological challenges to neonatal and juvenile ruminants is the development and establishment of the rumen. Using a subset of RNA-Seq data from our high-resolution atlas of gene expression in sheep (*Ovis aries*) we have provided the first comprehensive characterisation of transcription of the entire the gastrointestinal (GI) tract during the transition from pre-ruminant to ruminant. The dataset comprises 168 tissue samples from sheep at four different time points (birth, one week, 8 weeks and adult). Using network cluster analysis we illustrate how the complexity of the GI tract is reflected in tissue- and developmental stage-specific differences in gene expression. The most significant transcriptional differences between neonatal and adult sheep were observed in the rumen complex. Differences in transcription between neonatal and adult sheep were particularly evident in macrophage specific signatures indicating they might be driving the observed developmental stage-specific differences. Comparative analysis of gene expression in three GI tract tissues from age-matched sheep and goats revealed species-specific differences in genes involved in immunity and metabolism. This study improves our understanding of the transcriptomic mechanisms involved in the transition from pre-ruminant to ruminant. It highlights key genes involved in immunity, microbe recognition, metabolism and cellular differentiation in the GI tract. The results form a basis for future studies linking gene expression with microbial colonisation of the developing GI tract and will contribute towards identifying genes that underlie immunity in early development, which could be utilised to improve ruminant efficiency and productivity.

**Reference Numbers for Data in the Public Repositories:** The raw RNA-Sequencing data are deposited in the European Nucleotide Archive (ENA) under study accessions PRJEB19199 (sheep) and PRJEB23196 (goat). Metadata for all samples is deposited in the EBI BioSamples database under group identifiers SAMEG317052 (sheep) and SAMEG330351 (goat).

## Introduction

Sheep are an important source of meat, milk and fibre for the global livestock sector and belong to one of the most successful species of herbivorous mammals, the ruminants. Adult sheep have four specialized four specialized chambers comprising their stomach: fermentative fore-stomachs encompassing the rumen, reticulum and omasum and the “true stomach”, the abomasum (Dyce *et al.* 2010). The events surrounding the development of the rumen are among the most significant physiological challenges to young ruminants (Baldwin *et al.* 2004). As lambs transition from a milk diet to grass and dry pellet feed the gastrointestinal (GI) tract undergoes several major developmental changes. In neonatal lambs, feeding solely on milk, the fermentative fore-stomachs are not functional and the immature metabolic and digestive systems function similarly to that of a young monogastric mammal, with proteolytic digestion taking place inside the abomasum (Meale *et al.* 2017). At this stage the rumen has a smooth, stratified squamous epithelium with no prominent papillae (Baldwin *et al.* 2004). Suckling causes a reflex action that brings the walls of the reticulum together to form an ‘oesophageal’ or ‘reticular’ groove transferring milk and colostrum directly to the abomasum, where it is digested efficiently (Figure 1) (Dyce *et al.* 2010). In neonatal ruminants this is essential to ensure protective antibodies in the colostrum are transported intact to the abomasum.

**Figure 1:**
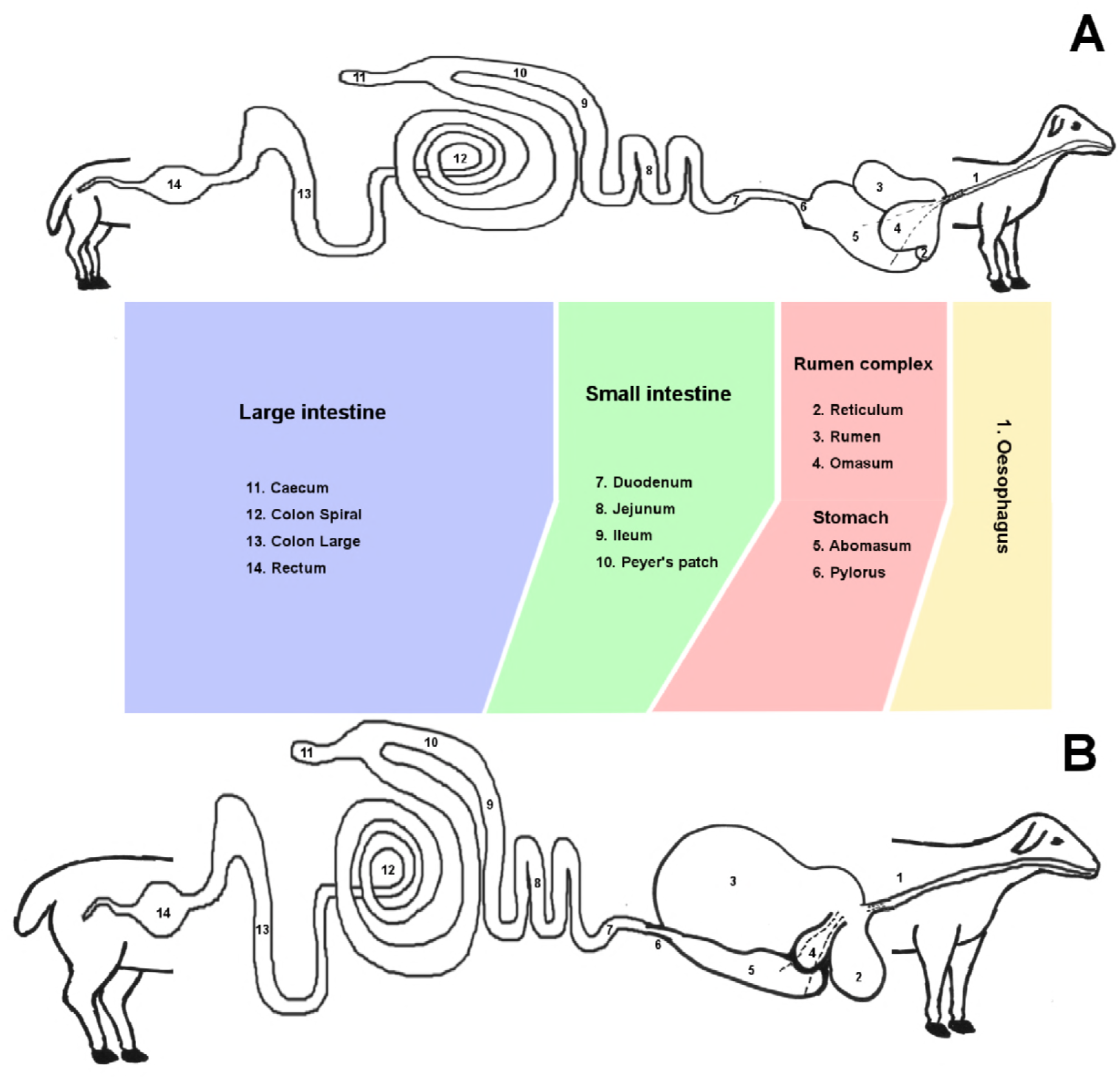
Diagrammatic representation of the morphological changes that occur in the gastrointestinal tract of a sheep during the transition from (A) pre-ruminant to (B) ruminant. The 14 regions sampled for this study are numbered. The oesophageal groove is indicated with dotted lines.

The introduction of grass and dry feed into the diet (which usually occurs in very small amounts from one week of age) inoculates the rumen with microbes. These proliferate, facilitating the digestion of complex carbohydrates which the adult ruminant relies upon to meet its metabolic needs, and stimulating growth and development of the rumen and reticulum (Bryant *et al.* 1958). The transition from pre-ruminant to ruminant occurs gradually from around 4 weeks of age. The rumen and reticulum are usually fully functional by the time the lamb reaches 8 weeks of age and has a completely grass and dry feed-based diet (Figure 1). The transition results in metabolic changes, as tissues shift from reliance on glucose supplied from milk to the metabolism of short-chain fatty acids as primary energy substrates. Whilst the most dramatic physical changes occurring during development are associated with the rumen epithelium, changes in intestinal mass, immunity and metabolism also occur in response to dietary changes (Baldwin *et al.* 2004).

Early development of the rumen complex and GI tract of sheep is of particular interest due to the transition from pre-ruminant to ruminant, which occurs over the period 8 weeks after birth. This period is crucial in the development of innate and acquired immunity, healthy rumen growth and establishment of the microbiome. These processes are likely to be intrinsically linked, as the GI tract protects the host from toxic or pathogenic luminal contents, while at the same time as supporting the absorption and metabolism of nutrients for growth and development (reviewed in (Steele *et al.* 2016; Meale *et al.* 2017)). Prior to the development of next generation sequencing technologies many studies used quantitative PCR to measure the expression of sets of candidate genes in ruminant GI tract tissues (reviewed in (Connor *et al.* 2010)). RNA-Sequencing (RNA-Seq) technology now provides a snapshot of the transcriptome in real-time to generate global gene expression profiles. This allows us to measure the expression of all protein coding genes throughout the development of the GI tract and associate these expression patterns with immunity, metabolism and other cellular processes at the gene/transcript level. The availability of high quality, highly contiguous, well annotated reference genomes for ruminant species, due in part to the efforts of the Functional Annotation of Animal Genomes Consortium (FAANG) (Andersson *et al.* 2015), has helped significantly in this task, particularly for sheep (Jiang *et al.* 2014) and goat (Bickhart *et al.* 2017; Worley 2017).

Previous studies characterising transcription in the GI tract throughout early development in ruminants examined links between feed intake and bacterial diversity and the development of the rumen (Connor *et al.* 2013; Wang *et al.* 2016; Xiang *et al.* 2016a). Another recent study characterised transcription in the adult rumen complex and GI tract of sheep, linking metabolic, epithelial and metabolic transcriptomic signatures (Xiang *et al.* 2016b). Similarly, Chao et al. 2017 performed transcriptional analysis of colon, caecum and duodenum from two breeds of sheep highlighting key genes involved in lipid metabolism (Chao *et al.* 2017). Others have linked diet to gene expression in the jejunal mucosa of young calves, suggesting that early feeding can have a profound effect on the expression of genes involved in metabolism and immunity (Hammon *et al.* 2018). The GI tract plays a significant role in defence against infection, because due to its large surface area it is often a site of primary infection via pathogen ingestion. This is particularly true in early development when the maturing lamb is exposed both to a range of pathogens from the environment and the rumen is colonised by commensal micro-organisms. Intestinal macrophages exhibit distinctive properties that reflect adaptation to a unique microenvironment (Bain & Mowat 2011). The microenvironment of the sheep GI tract changes dramatically during the transition from pre-ruminant to ruminant, and we hypothesise that the transcriptional signature of intestinal macrophages will reflect these physiological changes.

To characterise tissue specific transcription in the GI tract during early development we utilised a subset of RNA-Seq data, from Texel x Scottish Blackface (TxBF) lambs at birth, one week and 8 weeks of age and TxBF adult sheep, from our high resolution atlas of gene expression in sheep (Clark *et al.* 2017b). We characterise in detail the macrophage signature in GI tract tissues at each developmental stage and link these to other key biological processes ocurring as the lamb develops. We also perform comparative analysis of transcription in the rumen, ileum and colon of one-week old sheep with age-matched goats. One of the most significant physiological challenges to neonatal and juvenile ruminants is the development and establishment of the rumen. A clearer understanding of the transcriptomic complexity that occurs during the transition between pre-ruminant and ruminant will allow us to identify key genes underlying immunity in neonatal ruminants and healthy growth and development of the rumen and other tissues. These genes could then be utilised as novel targets for therapeutics in sheep and other ruminants and as a foundation for understanding rumen development, metabolism and microbial colonisation in young ruminants to improve efficiency and productivity.

## Materials and Methods

### Animals

Approval was obtained from The Roslin Institute and the University of Edinburgh Protocols and Ethics Committees. All animal work was carried out under the regulations of the Animals (Scientific Procedures) Act 1986. Full details of all the sheep used in this study are provided in the sheep gene expression atlas project manuscript (Clark *et al.* 2017b) and summarised in Table 1. GI tract tissues were collected from three male and three female adult Texel x Scottish Blackface (TxBF) sheep and nine Texel x Scottish Blackface lambs. Of these nine lambs, three were observed at parturition and euthanised immediately prior to their first feed. Three lambs were euthanised at one week of age pre-rumination (no grass was present in their GI tract) and three at 8 weeks of age once rumination was fully established. All the animals were fed *ad libitum* on a diet of hay and sheep concentrate nuts (16% dry matter without water), with the exception of the lambs pre-weaning (birth and one week of age) who suckled milk from their mothers. Goat GI tract and alveolar macrophage samples from one-week old un-weaned male goat kids were obtained from an abattoir (Table 1). We also included available RNA-Seq data from a trio of Texel sheep (a ewe, a ram and their female lamb) that was released with the sheep genome paper (Jiang *et al.* 2014). By incorporating this data in the analysis, we were able to include tissues not included in the TxBF samples e.g. omentum and an additional developmental stage (6-10 months) (Table 1).

**Table 1:**
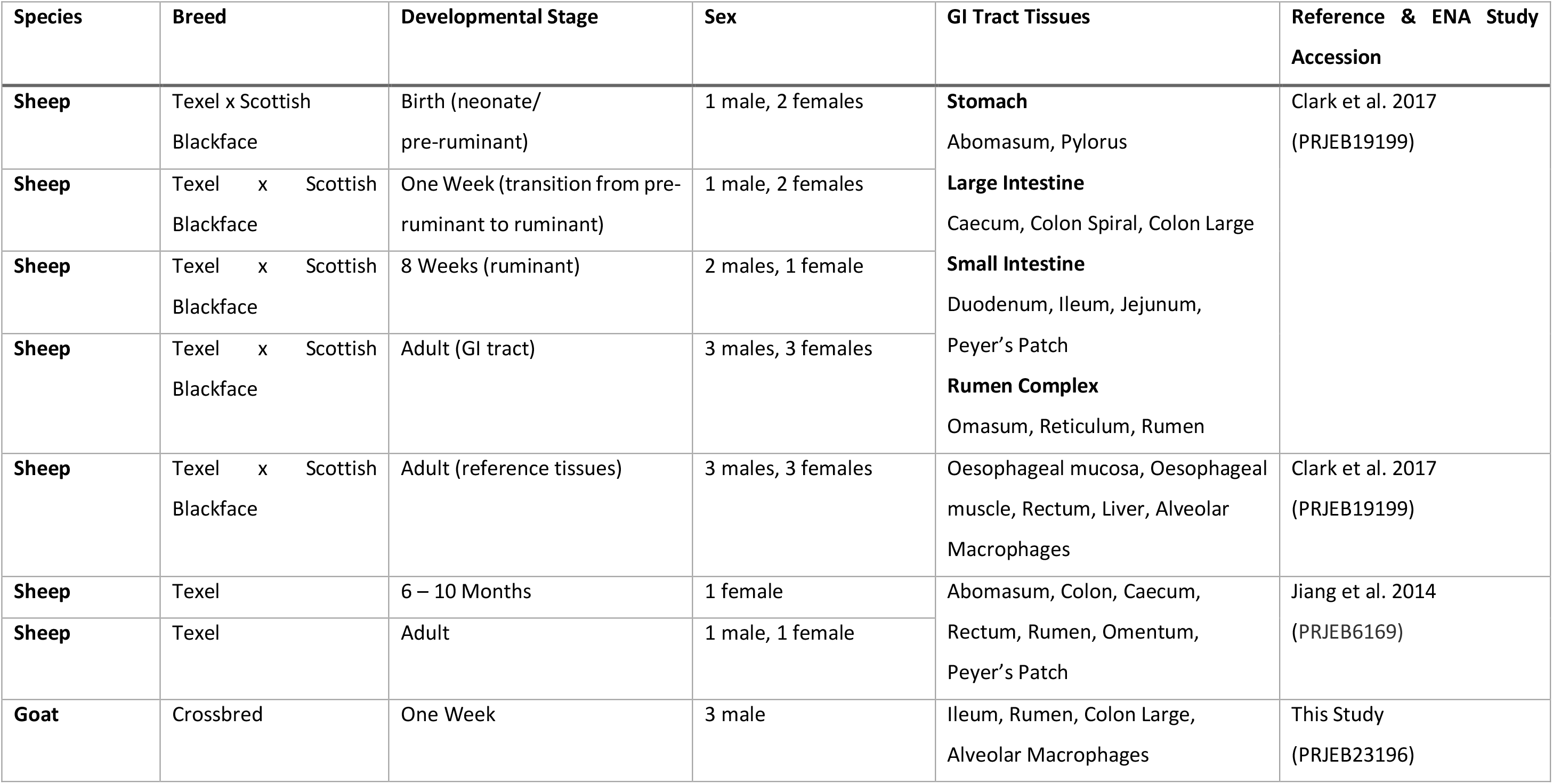
Details of animals and samples from the GI tract included in this study.

### Tissue Collection

In total thirteen different regions of the sheep GI tract were sampled, as detailed in Table 1 and illustrated in Figure 1. All post mortems were undertaken by the same veterinary anatomist and tissue collection from the GI tract was standardised as much as possible. All GI tract tissue samples were washed twice in room temperature sterile 1x PBS (Mg^2+^ Ca^2+^ free) (P5493; Sigma Aldrich) then chopped into small pieces <0.5cm and transferred to RNAlater preservation solution (AM7021; Thermo Fisher Scientific, Waltham, USA). To maintain RNA integrity, all GI tract tissue samples were harvested within 30 minutes from the time of death. Alveolar macrophages, liver, rectum and oesophageal muscle and tissue from the adult sheep were also included as reference tissues for comparison in the analysis. Isolation of alveolar macrophages from the adult sheep is described in detail in the sheep gene expression atlas project manuscript (Clark *et al.* 2017b). The goat samples (Table 1) were collected using the same methods as described for sheep.

### RNA Extraction and Library Preparation

We used a TRIzol^®^ (15596018; Thermo Fisher Scientific) based RNA extraction method which is described in detail in the sheep gene expression atlas project manuscript (Clark *et al.* 2017b). RNA quantity was measured using a Qubit RNA BR Assay kit (Q10210; Thermo Fisher Scientific) and RNA integrity estimated on an Agilent 2200 Tapestation System (Agilent Genomics, Santa Clara, USA) using the RNA Screentape (5067-5576; Agilent Genomics) to ensure RNA quality was of RIN^e^ > 7. RNA-Seq libraries were prepared by Edinburgh Genomics (Edinburgh Genomics, Edinburgh, UK) and run on the Illumina HiSeq 2500 (sheep) and Illumina HiSeq 4000 (goats) sequencing platform (Illumina, San Diego, USA). The GI tract tissues collected from the 9 TxBF lambs were sequenced at a depth of >25 million strand-specific 125bp paired-end reads per sample using the standard Illumina TruSeq mRNA library preparation protocol (poly-A selected) (Ilumina; Part: 15031047 Revision E). Depth refers to the number of paired end reads, therefore a depth of >25 million reads represents ~25M forward + ~25M reverse. The adult sheep GI tract tissues were also sequenced as above with the exception of the ileum, reticulum and liver which were sequenced using the Illumina TruSeq total RNA library preparation protocol (ribo-depleted) (Ilumina; Part: 15031048 Revision E) at a depth of >100 million reads per sample. The ileum and rumen samples from goats were sequenced at a depth of >30 million strand-specific 75p paired-end reads per sample using the standard Illumina TruSeq mRNA library preparation protocol (poly-A selected) (Ilumina; Part: 15031047 Revision E). The RNA-Seq libraries for the Texel dataset were also prepared by Edinburgh Genomics with RNA isolated using the method described in the sheep gene expression atlas project manuscript (Jiang *et al.* 2014; Clark *et al.* 2017b).

### Data Quality Control and Processing

The raw data for sheep, in the form of .fastq files, was previously released with the sheep gene expression atlas (Clark *et al.* 2017b) and is deposited in the European Nucleotide Archive (ENA) under study accession number PRJEB19199 (http://www.ebi.ac.uk/ena/data/view/PRJEB19199). The goat data is also deposited in the ENA under study accession number PRJEB23196 (http://www.ebi.ac.uk/ena/data/view/PRJEB23196). Both sets of data were submitted to the ENA with experimental metadata prepared according to the FAANG Consortium metadata and data sharing standards. Details of all the samples for sheep and goat, with associated data and metadata can also be found on the FAANG Data Portal (http://data.faang.org/) (FAANG 2017). The raw read data from the Texel samples incorporated into this dataset and previously published (Jiang *et al.* 2014) is located in the ENA under study accession PRJEB6169 (http://www.ebi.ac.uk/ena/data/view/PRJEB6169). The RNA-Seq data processing methodology and pipelines used for this study are described in detail in (Clark *et al.* 2017b). For each tissue a set of expression estimates, as transcripts per million (TPM), was obtained using the high-speed transcript quantification tool Kallisto v0.43.0 (Bray *et al.* 2016) with the Oar v3.1 reference transcriptome from Ensembl (Zerbino *et al.* 2018) as an index. Expression estimates for the GI tract dataset were then filtered to remove low intensity signals (TPM<1) and technical artefacts. To integrate expression estimates from the two different library types we performed a ratio correction of the TPM values as described in (Bush *et al.* 2017).

### Network Cluster Analysis

Network cluster analysis of the sheep GI tract dataset was performed using the network visualisation tool Graphia Professional (Kajeka Ltd, Edinburgh, UK) (Theocharidis *et al.* 2009; Livigni *et al.* 2018). To determine similarities between individual gene expression profiles a Pearson correlation matrix was calculated, for both sample-to-sample and gene-to-gene comparisons and filtered to remove relationships where *r* < 0.81 (sample-to-sample) and *r* < 0.85 (gene-to-gene). Network graphs were constructed by connecting nodes (genes or samples) with edges (where the correlation exceeded the threshold value). To interpret each graph a Markov Cluster algorithm (MCL) (van Dongen & Abreu-Goodger 2012) was applied at an inflation value (which determines cluster granularity) of 2.2. Visual examination was then used to interrogate the local structure of the graph. Similar samples/tissue types and genes with robust co-expression patterns formed clusters of highly interconnected nodes, implying related functions. To determine if genes within a cluster shared a similar biological function GO term enrichment based on gene ontology (Ashburner *et al.* 2000) was performed using the Bioconductor package ‘topGO’ (Alexa & Rahnenfuhrer 2010). For the gene-to-gene network analysis the top 20 largest clusters were assigned a functional class and sub-class based on GO term enrichment, gene function and the ‘guilt-by-association’ principle (Oliver 2000). Alveolar macrophage, oesophageal muscle and mucosa, rectum and liver from the adult sheep were included in the gene to gene network graph as ‘reference’ tissues for comparison, using a similar approach as described in (Xiang *et al.* 2016b). The Texel dataset (Jiang *et al.* 2014) was also included in the gene-to-gene network graph to add additional tissue samples and developmental stages not captured in the TxBF dataset. The sample-to-sample network analysis was used to illustrate transcriptional changes in tissues through each developmental stage (birth, one week, 8 weeks and adult) and included only the dataset from the TxBF sheep.

### Principal component analysis of transcriptional signatures in the developing GI tract

All statistical analysis was carried out in R (v >= 3.0.0) (R Core Team 2014) unless stated otherwise. We used Principal Component Analysis (PCA) to determine whether there was any strong age or tissue related transcriptional patterns observed in the GI tract. PCA was performed using FactoMineR v1.41 (Lê *et al.* 2008) with a subset of genes (n = 490) (extracted from the alveolar macrophage clusters (clusters 7 and 10) from the gene-to-gene network graph), centre scaled for computation of the principle components (PCs). The categorical data was excluded from the Eigen vector calculation (passed as qualitative variables). The top 5 and 10 PCs explained 62.6% and 76.9% of variability in the data respectively. In order to compare exploratory and discriminative power of PCs, the categorical data (age and tissue of origin) was overlaid on the PCs coordinate maps afterwards using centroid lines and colouring the observations by groups.

The expression levels as TPM of a small subset of macrophage and immune marker genes (*CD14, CD68, CD163* and *IL10*) were plotted as a HeatMap using ‘gplots’.

### Developmental Stage Specific Differential Expression Analysis for Sheep

Differential expression analysis was used to compare gene-level expression estimates from the Kallisto output as transcripts per million (TPM) across GI tract tissues and developmental time points, and between age-matched sheep and goats. The R/Bioconductor package tximport v1.0.3 was used to import and summarise the transcript-level abundance estimates from Kallisto for gene-level differential expression analysis using edgeR v3.14.0 (Robinson *et al.* 2010), as described in (Soneson *et al.* 2015; Love *et al.* 2017). For RNA-Seq experiments with less than 6 replicates per time point edgeR may be considered the optimal differential expression analysis package (Schurch *et al.* 2016). We selected three tissues (abomasum, rumen and ileum) as representative samples from 3 of the major compartments of the GI tract: the stomach, rumen complex and small intestine, respectively. Gene expression patterns in each tissue were compared between birth and one week, and one week and 8 weeks of age. To determine whether there was any tissue- and developmental stage-specific overlap of differentially expressed genes we generated Venn diagrams using the software tool Venny (Oliveros 2007). To investigate the function of the differentially expressed genes we performed GO term enrichment (Ashburner *et al.* 2000) using ‘topGO’ (Alexa & Rahnenfuhrer 2010). Only GO terms with 10 or more associated genes were included.

### Data Availability

All data analysed during this study are included in this published article and its additional files. The raw RNA-sequencing data are deposited in the European Nucleotide Archive (ENA) under study accessions PRJEB19199 (sheep) (https://www.ebi.ac.uk/ena/data/view/PRJEB19199) and PRJEB23196 (goat) (https://www.ebi.ac.uk/ena/data/view/PRJEB23196). Sheep data can also be viewed and downloaded via BioGPS (http://biogps.org/dataset/BDS_00015/sheep-atlas/) where the gene expression estimates for each tissue are searchable by gene name (http://biogps.org/sheepatlas). Sample metadata for all tissue and cell samples, prepared in accordance with FAANG consortium metadata standards, are deposited in the EBI BioSamples database under group identifiers SAMEG317052 (sheep) and SAMEG330351 (goat). All experimental protocols are available on the FAANG consortium website at http://www.ftp.faang.ebi.ac.uk/ftp/protocols. All supplementary material for this study has been deposited in Figshare.

## Results and Discussion

### Gene-to-gene network cluster analysis of the GI tract dataset

The dataset includes 168 RNA-Seq libraries in total from the TxBF described above sheep. Network cluster analysis of the GI tract data was performed using Graphia Professional (Kajeka Ltd, Edinburgh UK) (Theocharidis *et al.* 2009; Livigni *et al.* 2018). TPM estimates from Kallisto averaged across biological replicates (3 sheep per developmental stage) for the GI tract dataset were used to generate the network cluster graph. The full version of this averaged dataset was published with the sheep gene expression atlas and is available for download through the University of Edinburgh DataShare portal (http://dx.doi.org/10.7488/ds/2112). A version only including the TPM estimates for GI tract tissues, alveolar macrophages, thoracic oesophagus and liver is included here as Table S1.

The dataset was clustered using a Pearson correlation co-efficient threshold of *r* = 0.85 and MCL (Markov Cluster Algorithm (Gough *et al.* 2001)) inflation value of 2.2. The gene-to-gene network graph (Figure 2A) comprised 13,035 nodes (genes) and 696,618 edges (correlations above the threshold value). The network graph (Figure 2A) was highly structured comprising 349 clusters of varying size. Genes found in each cluster are listed in Table S2. Genes in Table S2 labelled ‘assigned to’ were annotated using an automated pipeline for the sheep gene expression atlas (Clark *et al.* 2017b). Clusters 1 to 20 (numbered in order of size; cluster 1 being the largest including 1724 genes in total) were annotated visually and assigned a broad functional ‘class’ (Table 2). Validation of functional classes was performed using GO term enrichment (Alexa & Rahnenfuhrer 2010) for molecular function, cellular component and biological process (Table S3). Figure 2B shows the network graph with the nodes collapsed by class, and the largest clusters numbered 1 to 20, to illustrate the relative number of genes proportional to the size of each cluster.

**Figure 2.**
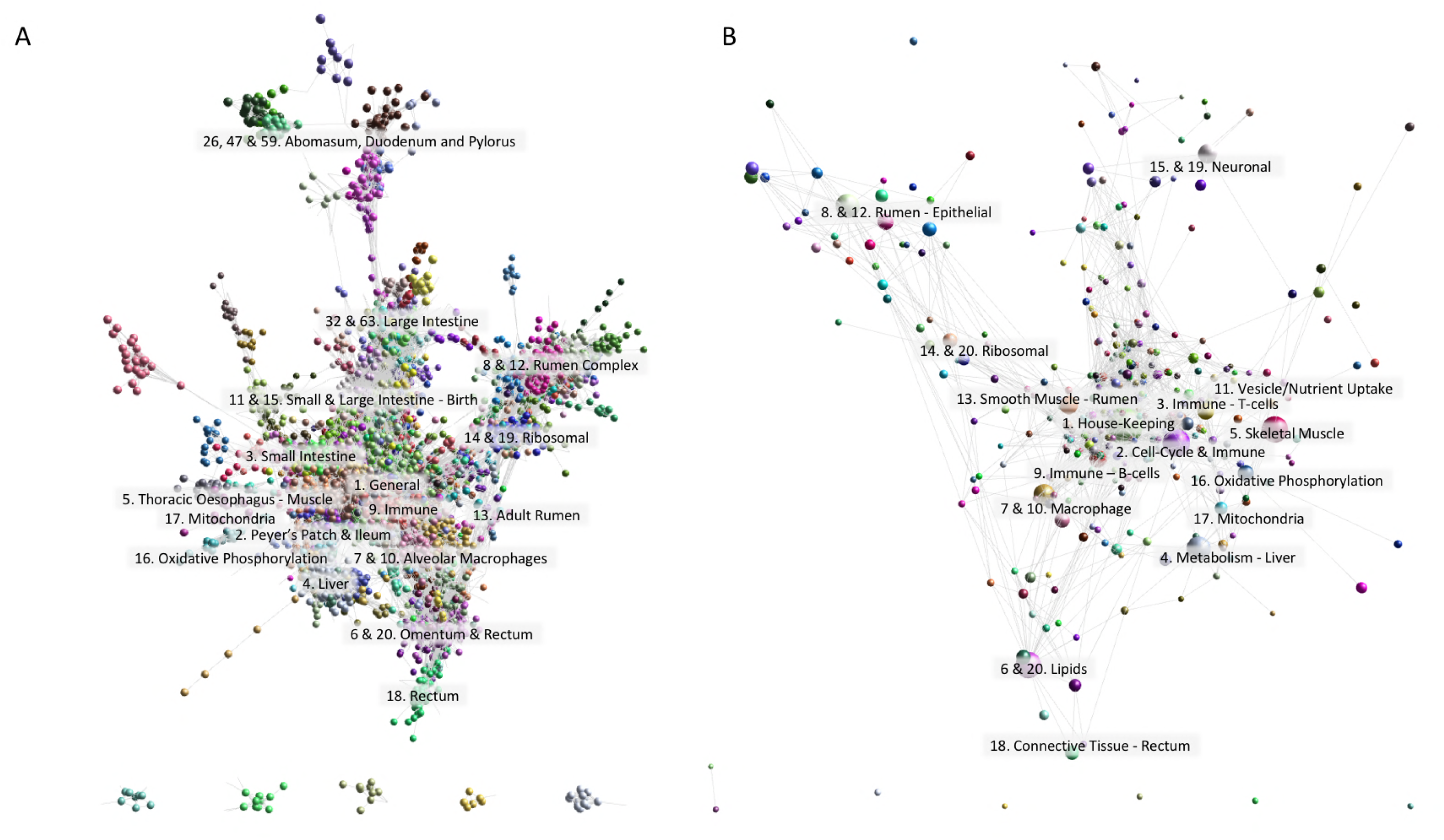
A: Gene-to-gene network graph of sheep GI tract tissues and including alveolar macrophages, oesophageal tissue and liver. The top 20 largest clusters are annotated by functional class. B: Gene-to-gene network graph with the nodes collapsed by class to illustrate the relative size of each cluster. Created using Graphia Professional with parameters Pearson’s R=0.85, MCLi=2.2, Minimum Component Size=2, Minimum Cluster Size=2.

**Table 2:** Functional annotation of the top 20 clusters for the sheep GI tract dataset including liver, oesophagus and alveolar macrophages as reference samples.

| Cluster | Functional Class | Tissue Specific Expression | Marker Genes |
| --- | --- | --- | --- |
| 1 | House-Keeping (1) | Ubiquitous | *ATM, CDK9, CEP63* |
| 2 | Cell-Cycle & Immune | Peyer's Patch, Ileum | *CBX2, CD19, CDK1, CKLF* |
| 3 | Immune (T-Cells) | Small and Large Intestine | *CD4, CD8A, IL16, IL2RA, PARP8* |
| 4 | Metabolism | Liver | *SERPINA10, SLC10A1, FGF12, IGF1, IGF2* |
| 5 | Muscle - skeletal muscle | Thoracic Oesophagus | *MYH4, MYL2, OXCT1, ACTA1* |
| 6 | Lipids | Omentum, Rectum | *PLIN1, ADIPOQ* |
| 7 | Immune - Macrophages | Alveolar Macrophages | *CLEC4E, CD68, CCL4, CSF1, TLR4, CCR1, ITGAX, ITGAM, ADGRE1, MERTK, ICAM-1* |
| 8 | Epithelial - Rumen | Rumen Complex - General | *KRT15, N4BP3, NOD1, SOX2* |
| 9 | Immune | Ileum, Peyer's Patch, Alveolar Macrophages | *IL10, CD73, CLEC10A* |
| 10 | Immune - Macrophages | Alveolar Macrophages | *CD83, CD44, TNF-alpha, VSIG4, P2RY12, JUNB* |
| 11 | Vesicle Formation/Nutrient Uptake | Small Intestine (at birth) | *SLC10A4, SLC16A11, AQP7, AQP8* |
| 12 | Epithelial/Immune - Rumen | Rumen Complex (increasing with age) | *KRT5, PIP, IL19, VSIG8, IL36A, IL36B* |
| 13 | Smooth Muscle | Adult Reticulum and Rumen | *FIBIN, MYL, FMOD, CALD1, MYH11, ACTG2, CNN1* |
| 14 | Ribosomal | Ubiquitous | *RPS3, RPL32* |
| 15 | Neuronal | Large and small intestine (up-regulation at birth) | *IGF2BP1, MAPK10, FBXL16, VIP* |
| 16 | Oxidative Phosphorylation | Ubiquitous | *ATP5B, COX10, NDUFA3* |
| 17 | Mitochondria | Ubiquitous | *PLAG1, F3, ESR2* |
| 18 | Connective Tissue | Rectum | *COL6A6, GDF6* |
| 19 | Neuronal | True stomach, large and small intestine | *CPS1, DEPCR2* |
| 20 | Ribosomal | Ubiquitous | *RPL5, RPL15* |

### Complexity of cell types within the GI tract is reflected in gene-to-gene network clustering

The GI tract is a highly complex organ system in ruminants with region-specific cellular composition. This complexity is illustrated by the highly structured gene-to-gene network graph of GI tract tissues (Figure 2A). Other studies have characterised in detail the transcriptional signatures in the GI tract of adult sheep (Xiang *et al.* 2016a; Xiang *et al.* 2016b) and as such we will only broadly describe the clusters here and focus instead on developmental stage-specific transcriptional patterns. As is typical of large gene expression datasets from multiple tissues cluster 1, the largest cluster, was comprised largely of ubiquitously expressed ‘house-keeping’ genes, encoding proteins that are functional in all cell types. Enriched GO terms for genes within this cluster were for general cellular processes and molecular functions performed by all cells including RNA-processing (p=<1×10^−30^) and histone binding (p=3.9×10^−8^) (Table S3).

Although cluster 1 exhibited a general expression pattern the majority of the rest of the clusters (with the exception of cluster 2, discussed below) exhibited some tissue/cellular process specific gene expression pattern. The majority of clusters included genes associated with more than one cell type/cellular process. This is because the lamina propria, one of the three layers of the mucosa, which lies beneath the epithelium and lines the majority of the GI tract, is made up of different cell types including endothelial, immune and connective tissues (Takahashi-Iwanaga & Fujita 1985). As a consequence expression signatures of components of the lamina propria were common across the network graph for sheep, as previously reported from transcriptional analysis of the pig GI tract (Freeman *et al.* 2012).

The GI tract of the sheep is lined with epithelium, and the rumen in particular exhibits a strong epithelial signature, which differentiates it transcriptionally from other tissues (Xiang *et al.* 2016a; Xiang *et al.* 2016b). Genes in clusters 8 and 12, expressed in the rumen, exhibited a typical epithelial signature, and included genes in the KRT superfamily e.g. *KRT15*, with enriched GO terms for keratinocyte differentiation (p=1.4×10^−7^ and p=6.8×10^−7^, respectively). The rumen papillae help to increase the surface area available for the absorption of nutrients. The small intestine is also a major site of absorption and transport of nutrients. One small intestine specific cluster (cluster 11) was comprised of genes involved both in nutrient uptake and vesicle formation. Cluster 11 contained numerous genes in the solute carrier super family, a group of membrane transport proteins including *SLC10A4* and *SLC16A11*. Genes encoding water channel proteins were also found in cluster 11 including aquaporins *AQP7* and *AQP8*. Significant GO terms for cluster 11 included acrosomal vesicle (p=0.0033) and peptidyl-tyrosine autophosphorylation (p=0.00209).

Genes in cluster 5 were expressed in thoracic oesophageal skeletal muscle tissue and associated with muscle fibre development (p=1.3×10^−10^) and skeletal muscle tissue development (p=4.4×10^−12^). Several skeletal muscle specific genes were found within this cluster including, *ACTA1*, which is associated with skeletal muscle function and encodes a product belonging to the actin family of proteins (Clarke et al. 2007). Similarly, cluster 13 included genes associated with structural constituents of muscle (p=0.00108) but in this case the smooth muscle that surrounds the GI tract. Genes in cluster 13 were predominantly expressed in the rumen complex of adult and 8 week old sheep, and included several genes related to smooth muscle function and regulation (Table 2). *CALD1* for example encodes a calmodulin- and actin-binding protein that plays an essential role in the regulation of smooth muscle and nonmuscle contraction (Huang et al. 2010). Intestinal motility, is the result of contraction and relaxation of the smooth muscle, and a key function of the GI tract.

The numerous functions of the GI tract such as intestinal motility, exocrine and endocrine secretion, immune function and circulation are under complex neural control. Clusters 15 and 19 were associated with neuronal cells and included GO terms for synapse (p=0.000018) and synaptic membrane (p=4.9×10^−7^). However, overall expression of these genes was low relative to other clusters, presumably because neurons comprise only a small percentage of the cells that make up the GI tract, and therefore their expression level would be reduced compared to other more abundant cell types. A similar trend was observed in analysis of neuronal signatures in the pig GI tract (Freeman *et al.* 2012). This might also account for an increased expression level at birth of genes in cluster 15 as there will be comparatively fewer other cell types differentiated at this time point.

Several clusters in the top 20 largest clusters exhibited a strong immune signature and are discussed in more detail below. Analysis of the gene-to-gene network graph revealed the GI tract of the sheep comprises at least five major cell types: epithelia, immune cells, neuronal cells and mesenchymal cells (muscle, connective tissue). Interrogation of the expression profiles for each cluster in the gene-to-gene network graph revealed that cell type and tissue specific expression signatures varied with developmental stage (discussed in more detail below).

### Strong immune signatures highlight the role of the GI tract as an immune organ

The GI tract has the largest surface area in the body in contact with the external environment. The intestinal epithelium, a single-cell layer, is the barrier between the external environment of the lumen (which contains pathogens, antigens and commensal bacteria) and the body. As such it is not surprising that the second largest cluster of genes (cluster 2) contained many genes associated with the immune response, their expression being two- to three-fold higher in the ileum and Peyer’s patches than other regions of the GI tract. This pattern of expression was also observed in pig (Freeman *et al.* 2012). The lower small intestine Peyer’s patches (small masses of lymphatic tissue found throughout the ileum) form an important part of the immune system by monitoring intestinal bacteria populations and preventing the growth of pathogenic bacteria (Jung *et al.* 2010). Included in cluster 2 were genes encoding many of the protein components of the B cell receptor complex (*CD19*, *CD79B*, *CR2*) (Treanor 2012). Also evident in this cluster were many genes associated with the cell cycle, for example cyclins, DNA polymerases and kinesins, which were identified in the sheep gene expression atlas as a cell cycle specific cluster (Clark *et al.* 2017b). Significant GO terms for cluster 2 include G2/M transition of mitotic cell cycle (p=1.8×10^−9^) and meiotic cell cycle process (p=6.9×10^−9^). The high level of lymphocyte proliferation and replenishment of intestinal macrophages by a high turnover of monocytes (Bain & Mowat 2011; Shaw *et al.* 2018), and therefore the frequency of cells undergoing mitosis in the Peyer’s patches and Ileum, might explain the association of cell cycle genes with an immune signature (David *et al.* 2003).

Other GI clusters with an immune signature included clusters 3, 7, 9 and 10. Cluster 3 exhibited a strong immune signature, particularly associated with T-cells, with significant GO terms for positive regulation of T-cell activation (p=2.1×10^−21^) and cytokine receptor activity (p=5.4×10^−5^). Genes within cluster 3 included the T-cell marker genes *CD3, CD4* and *CD6,* as well as Toll-like receptor genes *TLR1* and *TLR10* (Akira & Takeda 2004). Expression was highest for this cluster in the small intestine particularly the ileum and Peyer’s patches. Cluster 9 exhibited a similar immune related expression pattern including genes involved in T-cell-B-cell interactions such as *CD37*, and cytokine production such as *IL10* and *CD80*. Significant GO terms for cluster 9 included B cell receptor signalling pathway (p=4.9×10^−6^) and regulation of B-cell activation (p=6.3×10^−6^) as well as Toll-like receptor 4 signalling pathway (p=2.8×10^−5^).

Macrophages play an essential role in inflammation and protective immunity and in the GI tract are important for local homeostasis, maintaining a balance between microbiota and the host immune system (Mowat & Bain 2011). Clusters 7 and 10 contained many macrophage specific marker genes including *CD68* and *ADGRE1 (EMR1)* with enriched GO terms for the immune response (p=7.3×10^−6^). Many of these genes are also commonly used as marker genes for intestinal macrophages including, *CCRL1* (synonym *CX3CR1*), *ITGAX* (synonym *CD11C*) and *ITGAM* (synonym *CD11B*) (Mowat and Bain 2011; Bain and Mowat 2011; Shaw et al. 2018). Other macrophage marker genes such as *CD14* and *CD163* were found in a much smaller macrophage-specific cluster, cluster 154, indicating some tissue specific differences in macrophage expression.

We included alveolar macrophages in the gene-to-gene network clustering of the GI tract dataset to look specifically at the expression of C-type lectins, and other receptors involved in bacterial recognition, in the GI tract relative to their expression in other populations of macrophages. Upon initial encounter with pathogens, the immune system needs to rapidly recognise these as potentially harmful. Innate immune cells, such as macrophages, use a limited number of pattern recognition receptors (PRRs), including C-type lectin receptors (CLRs), which activate immediate anti-microbial effectors or other mechanisms for defence against pathogens (reviewed in (Akira & Takeda 2004)). Analysis of the pig gene expression atlas revealed that there were a large number of genes for C-type lectins that were highly-expressed in alveolar macrophages but appeared down-regulated in the pig GI tract (Freeman *et al.* 2012). To determine if this was also the case in sheep we examined the expression of six C-type lectin marker genes (*CD68*, *CLEC4D*, *CLEC4E, CLEC5A*, *CLEC7A* and *SIGLEC1*) in sheep using the gene expression profiles for the sheep atlas dataset on BioGPS (Clark *et al.* 2017a). In sheep, as in pig, the four C-type lectins and *SIGLEC1* all exhibited very weak expression in the GI tract tissues. *CLEC4D, CD68*, *CLEC4E* and *CLEC7A* were all highly expressed in sheep alveolar macrophages, as was observed in pig. *CLEC5A* and *SIGLEC1* did not show the same upregulation in sheep alveolar macrophages as they did in pigs indicating these genes exhibit a more species-specific expression pattern in alveolar macrophages. To investigate whether these trends also applied to other ruminants, we compared the expression of these genes in alveolar macrophages, ileum, large colon and rumen from one-week old sheep with age-matched goats (Figure 3). Similar expression patterns were observed for goat as for sheep, the only exception being *CLEC5A* which was upregulated in goat alveolar macrophages, unlike in sheep. Generally, the high expression of gene for C-type lectins in alveolar macrophages and down regulation in the GI tract appears to be conserved between both ruminants and monogastric mammals.

**Figure 3:**
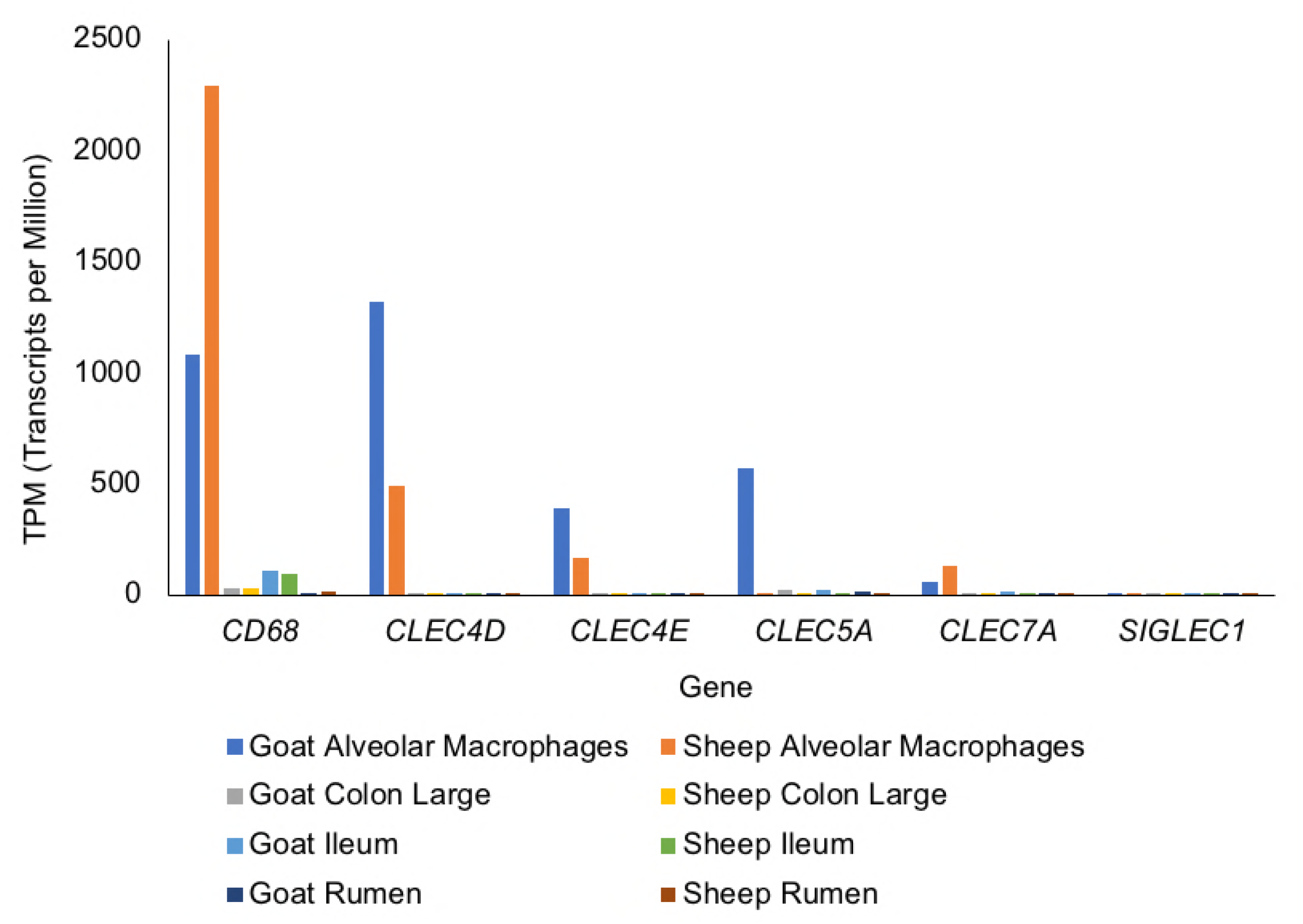
Comparative analysis of the expression of five C-type lectin genes, measured as transcripts per million (TPM), across three GI tract tissues (Large Colon, Ileum and Rumen) and alveolar macrophages in sheep and goats.

The expression pattern of *CD68, CLEC4D, CLEC4E* and *CLEC7A*, which are upregulated in alveolar macrophages and down regulated in the GI tract, indicate they are likely to be necessary for the elimination of inhaled pathogens, where such inflammatory responses to microbes would be undesirable in the intestine. *CLEC4E*, for example, has been shown to be important in protective immunity against pneumococcal pneumonia in lung macrophages in mice (Behler-Janbeck *et al.* 2016) and *CLEC7A* is involved in the innate immune respiratory response to fungal pathogens (Saijo & Iwakura 2011). Although ruminants have a larger and more diverse microflora in the GI tract compared to pigs, which are monogastric mammals (O’Donnell *et al.* 2017), expression patterns of the C-type lectin marker genes appear to be similar between the two species. Our findings generally support those observed in pig that intestinal macrophages differ transcriptionally from lung and blood macrophage populations, which might be because they have adapted to be hypo-responsive to food-derived glycoproteins while alveolar macrophages need to utilise the receptors to both recognize and engulf potential pathogens (Freeman *et al.* 2012).

### Network cluster analysis and principal component analysis of tissue samples reveals a strong effect of developmental stage on tissue specific transcription

Sample-to-sample network cluster analysis was used to visualise the effect of developmental stage on tissue specific gene expression. To perform the sample-to-sample network cluster analysis we used a version of the GI tract dataset that was not averaged across individuals and did not include the Texel dataset or the alveolar macrophage (AM), oesophagus and liver samples (Figure 4 A&B). The full version of this dataset was published with the sheep gene expression atlas and is available for download through the University of Edinburgh DataShare portal (http://dx.doi.org/10.7488/ds/2112). A version only including the TPM estimates used for this analysis is included here as Table S4.

**Figure 4:**
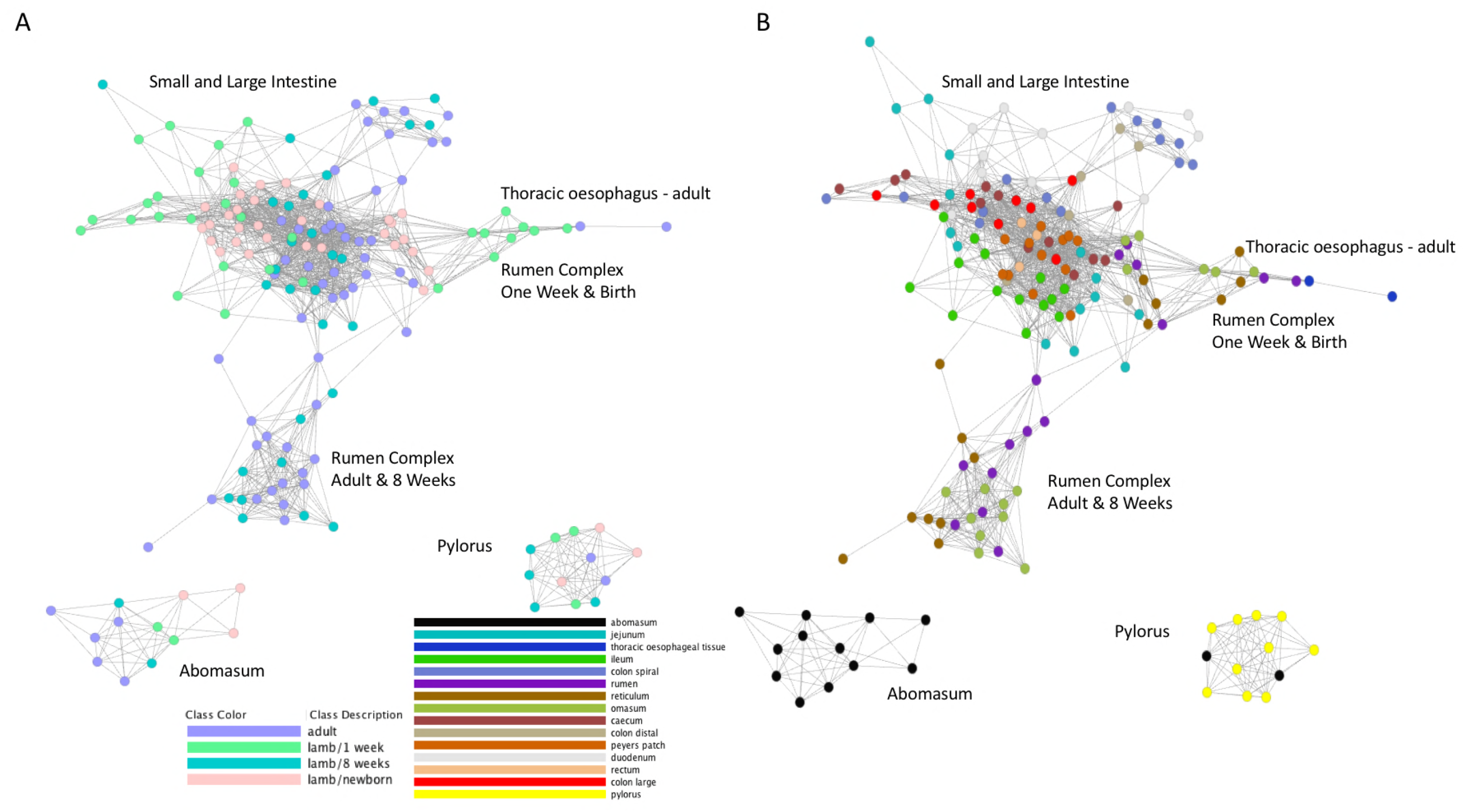
Sample-to-sample network graph of sheep GI tract samples coloured according to developmental stage (A) and tissue (B). Created using Graphia Professional with parameters Pearson’s R=0.85, MCLi=2.2, Minimum Component Size=2, Minimum Cluster Size=2.

Developmental stage specific transcriptional patterns were reflected particularly strongly in the rumen complex (reticulum, rumen and omasum) tissue samples (Figure 4A). Rumen complex samples from lambs at birth and one week of age formed a distinct cluster from rumen complex samples at 8 weeks of age and adult. This reflects the changes in tissue structure, function and morphology of the reticulum, rumen and omasum in the transition from pre-ruminant to ruminant (Figure 1). The close proximity of the adult oesophageal samples and the pre-ruminant rumen complex samples in the network cluster analysis indicate that tissue from the rumen complex at birth and one week of age is transcriptionally similar to adult oesophageal tissue in comparison with tissue from the rumen of adult sheep (Figure 4A). At birth and one week of age the walls of the reticulum join together to form an ‘oesophageal’ or ‘reticular’ groove transferring milk and colostrum directly to the abomasum, where it is digested most efficiently bypassing the rumen (Figure 1) (Meale *et al.* 2017). Rumen tissue of lambs at birth and at one week of age will therefore be functionally and transcriptionally different to rumen tissue of adult sheep. The rumen of neonatal lambs, for example, is essentially non-functional in ketogenic capacity and does not exhibit the same high degree of keratinisation that is characteristic of the adult rumen (reviewed in (Baldwin *et al.* 2004)). Consequently, the strong epithelial and metabolic signatures of the adult rumen (Xiang *et al.* 2016b) are likely to be weaker at one week and birth driving some of the observed cluster separation between the developmental stages (Figure 4A).

The sample-to-sample graph also illustrates the strong tissue specific differences in transcription (Figure 4B). The pylorus and abomasum samples, for example, form distinct clusters from the other regions, reflecting the fact that they have unique functions. The pylorus and abomasum both have a glandular lining, with various glandular cell types involved with enzyme secretion, lubrication, and endocrine control and specialized structures, such as the pyloric and fundic glands of the stomach (Dyce *et al.* 2010). The large and small intestine and rumen complex also form large separate clusters highlighting the transcriptional similarities between tissues within and between these regions of the GI tract (Figure 4B).

Developmental stage- and tissue-specific transcriptional patterns observed in the sample to sample graph were also reflected in principal component analysis (PCA) of the dataset (Figure S1). Tissue specific clusters of samples were largely separated by organ system (rumen complex, small and large intestine, oesophagus and stomach) (Fig S1 A&B). The effect of age on transcription was particularly evident in PC2, with samples from newborn lambs clustered separately from the other 3 time points (Figure S1 C&D).

### Developmental stage specific changes in immune gene signatures occur during the transition from pre-ruminant to ruminant

One of the largest macrophage populations in the body occupies the lamina propria of the GI tract (Bain & Mowat 2011). Macrophages are so numerous that the expression of macrophage-related genes can be detected within the total mRNA from GI tract samples. Constant exposure to food and environmental antigens and a wide diversity of commensal bacteria make the GI tract of developing sheep a primary site for pathogen infection (Mantovani & Marchesi 2014). As a consequence, GI immune cells are involved in a balanced immune response, focussed on controlling pathogen invasion while recognising commensal colonising microbes. This response will change as the lambs age reflecting changes in diet (the transition from milk to pellets and hay) and environment (transition from maternal transmission of pathogens to those from the surrounding environment). Intestinal mononuclear phagocytes, of which monocyte-derived macrophages form the most abundant population play a key role in the maintenance of the intestinal immune response to pathogens and homeostasis (Mantovani & Marchesi 2014). Macrophage specific genes are therefore likely to be driving some of the observed transcriptional differences observed between lambs at birth and one week of age and 8 week old and adult sheep, described in the previous section (See Figure 4 and Figure S1).

We used PCA to examine the clustering pattern of a set of 490 macrophage associated genes (extracted from the alveolar macrophage specific clusters 7 and 10 from the gene to gene network graph, detailed in Table S2) (Figure 5). We looked at tissue specific transcriptional patterns (Figure 5A) and found the small intestine and rumen complex distinguished by components PC1 and PC2, respectively, indicating differences in the macrophage expression profile in these tissues. A similar effect was observed between components PC1 and PC3 (Figure 5B). Developmental stage specific transcriptional patterns in macrophage specific genes were evident when comparing components PC1 and PC3 (Figure 5C), and particularly when comparing PC3 and PC4 (Figure 5D), where the samples from newborn lambs are clearly distinct from the other developmental stages. These differences perhaps reflect that pathogen challenge in the newborn lambs will be reduced immediately at birth and therefore the inflammatory response would be less than at later developmental stages. In addition, the newborn lambs would not have received any colostrum which is key in the homologous transfer of passive immunity between the mother and neonate, and the initial source of acquired immunity by the newborn lamb (Stelwagen *et al.* 2009). A wide variety of immune components linked to the innate immune response have been identified in colostrum and milk including neutrophils and macrophages as well immunomodulatory factors (including numerous pro- and anti-inflammatory cytokines) and peptides and proteins with direct antimicrobial activity (Stelwagen *et al.* 2009). Another potential driver of these differences is that the GI tract will gradually become colonised with gut macrophages, and at birth there will be a comparatively higher proportion of monocytes (Shaw *et al.* 2018). The GI tract harbours multiple distinct populations of macrophages, monocytes and other immune cells that exhibit some level of stochasticity over time (Bain & Mowat 2011; Shaw *et al.* 2018) which is reflected in the observed transcriptional patterns. The macrophage populations that are most numerous in neonatal lambs may be quite different from those that are prevalent in the GI tract of adult sheep and this will vary according to tissue.

**Figure 5:**
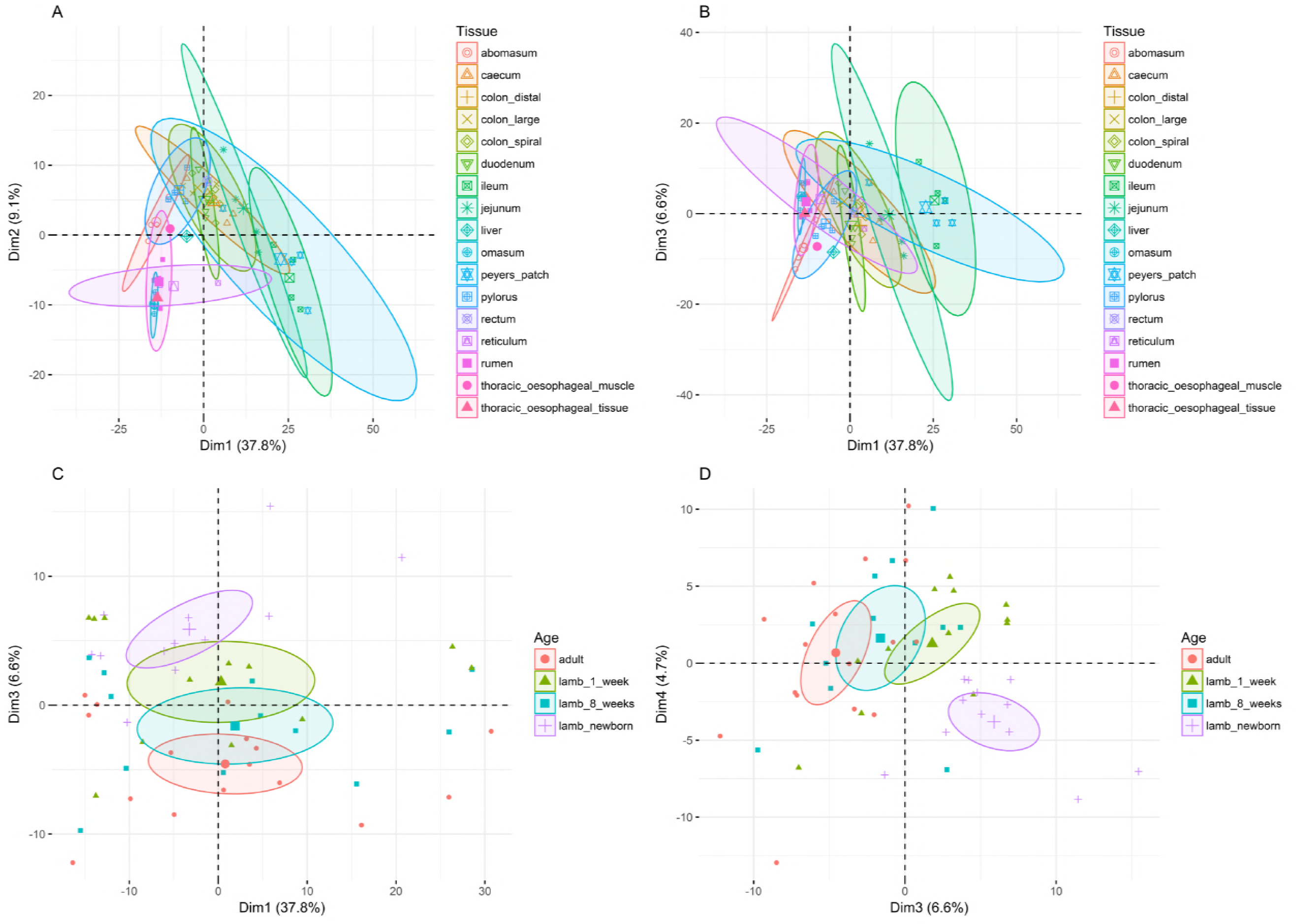
Principal Component Analysis of macrophage specific signatures in the sheep GI tract illustrating separation by tissue (A & B) and developmental stage (C & D) in three and four components, respectively.

To look in more detail at developmental stage specific differences in expression of monocyte and macrophage marker genes we compared the expression of *CD14, CD68, CD163* and *IL10* across GI tract tissues at each developmental stage (including liver and alveolar macrophage samples for reference) (Figure 6). *CD14, CD68* and *CD163* were upregulated in alveolar macrophages as expected (Gordon *et al.* 2014). This is not immediately obvious form the heatmap however due to high level of expression of CD68 (2291 TPM) relative to the levels of CD14 (171 TPM) and CD163 (168 TPM). Expression of *CD68* was highest in the ileum relative to other GI tract tissues, which probably reflects the fact that ileum contains the largest proportion of tissue macrophages relative to the other GI tract tissues sampled. The expression of the monocyte marker gene *CD14* increased as the lambs aged across most of the GI tract tissues sampled. This may reflect the lower number of monocytes in neonatal lambs relative to juvenile and adult animals (Kramer *et al.* 2003) or the high turnover of blood monocytes and continual replenishment of intestinal macrophages by monocytes that has been reported in the GI tract (Bain *et al.* 2016). The effect was most obvious in the rumen complex tissues in which *CD14* expression increased with development of the rumen. In contrast *CD163* and *CD68* expression decreased as the lambs aged in the rumen complex but increased in the developing small and large intestine. These differences in expression of macrophage marker genes reflect tissue specific differences in macrophage colonisation with age and the changing function of the rumen through these developmental stages. *CD163*, for example, directly induces intracellular signaling leading to secretion of anti-inflammatory cytokines (Moestrup & Moller 2004), and has been shown to be a macrophage receptor for bacteria (Fabriek *et al.* 2009). During bacterial infection *CD163* on resident tissue macrophages acts as an innate immune sensor, inducing local inflammation (Fabriek *et al.* 2009). Induction of *CD163* might therefore be less desirable in the rumen once colonisation with intestinal microbes has been established, accounting for the decrease in expression of *CD163* in the rumen complex. The increase in expression of *CD163* in the ileum and Peyer’s patches with age reflects the gradual colonisation of these tissues with intestinal macrophages and their importance in the innate immune response.

**Figure 6:**
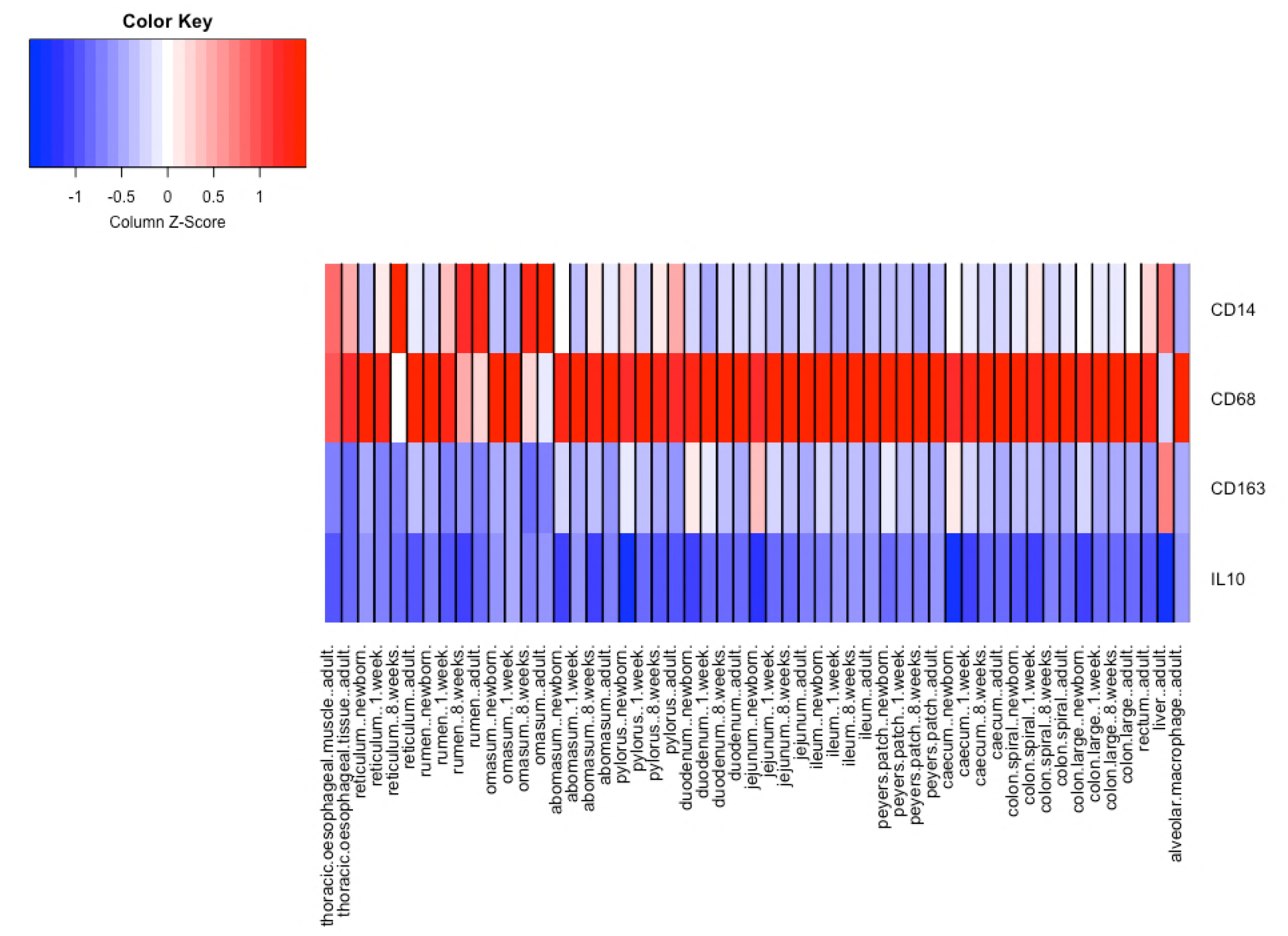
Red/blue heat map visualisation of the expression of a set of four immune marker genes (*CD14, CD68, CD163* and *IL10*) across sheep GI tract tissues at each developmental stage (including thoracic oesophagus, liver and alveolar macrophage samples for reference). Red indicates up-regulation of genes and blue down-regulation using z-score normalisation of TPM values for each gene.

The expression of *IL10* was very low across all the GI tract tissues which is in contrast to other mammalian species where it has been shown to be constitutively expressed in intestinal macrophages, reviewed in (Bain & Mowat 2011). For the majority of tissues TPM was <3 with the exception of the ileum and Peyer’s patch where expression ranged between 6 and 12 TPM. Interleukin 10 (*IL10*) is a key anti-inflammatory cytokine that can inhibit proinflammatory responses of both innate and adaptive immune cells (Mantovani & Marchesi 2014; Shouval *et al.* 2014). A relative reduction in *IL10* expression in the GI tract could indicate a suppressed macrophage response to intestinal microbes. This could be of potential benefit for ruminant species, which have a different relationship with intestinal flora than monogastric mammals and this would be interesting to explore further.

### Differential expression analysis reveals little overlap between stage-specific transcriptional signatures

To characterise which genes were driving transcriptional patterns, more broadly, during the transition from pre-ruminant to ruminant we chose three regions of the GI tract (rumen complex, stomach and small intestine) and selected one tissue per region (rumen, abomasum and Ileum) to perform differential expression analysis using pairwise comparisons between birth and one week and one week and 8 weeks of age (Table S5 – Rumen, Table S6 – Abomasum and Table S7 – Ileum). Differentially expressed genes are detailed in S5 Table (rumen), S6 Table (abomasum) and S7 Table (ileum); those of particular interest are shown in Table 3. To determine the main functions of genes exhibiting differential expression in each comparison we used GO term enrichment (Table S8). GO terms related to immunity were common across all three tissues but particularly the rumen and ileum between birth and one week of age. In the ileum between birth and one week of age enriched GO terms were predominantly associated with immunity and included ‘immune response’, ‘chemokine activity’, ‘chemokine production involved in the inflammatory response’ as well as others related to metabolism. In contrast between one and 8 weeks of age GO terms were associated predominantly with metabolic processes, and vesicle formation. A similar but less exaggerated trend was also observed in the rumen, with GO terms for ‘defense response to other organisms’, ‘immune response’ and more generally for metabolism enriched between birth and one week of age and then a shift towards metabolic and muscle and epithelial differentiation between one and 8 weeks. In the abomasum enriched GO terms between both sets of time points were more similar than for the other two tissues which probably reflects the fact that the functional changes in the abomasum are comparatively less than the other two tissues throughout the transition from pre-ruminant to ruminant.

**Table 3:** Examples of differentially expressed genes identified using pairwise comparison of abomasum, rumen and ileum samples from sheep at birth vs one week and one-week vs 8 weeks.

| Gene | Tissue | Developmental Stage | Role | Fold Change | P-Value | Direction | TPM 1 | TPM 2 |
| --- | --- | --- | --- | --- | --- | --- | --- | --- |
| **PIP** | Rumen | 1 vs 8 Weeks | Mucosal Immunity | 3.1 | 5.1x10<sup>-7</sup> | Up | 375 | 2113 |
| **CLEC4E** | Ileum | 1 vs 8 Weeks | Inflammatory and Immune Response | 2.4 | 2.7x10<sup>-4</sup> | Up | 15 | 76 |
| **IL36A** | Rumen | 1 vs 8 Weeks | Inflammatory Response | 3.7 | 1.4x10<sup>-5</sup> | Up | 275 | 2647 |
| **JAK3** | Rumen | 0 vs 1 Week | Cytokine signalling | 5.5 | 2.1x10<sup>-83</sup> | Up | 115 | 4311 |
| **ISG15** | Abomasum | 1 vs 8 Weeks | IFN response | 3.1 | 1.1x10<sup>-67</sup> | Up | 524 | 602 |
|  | Abomasum | 0 vs 1 Week | IFN response | 2.6 | 1.2x10<sup>-36</sup> | Up | 15 | 99 |
|  | Rumen | 0 vs 1 Week | IFN response | 6.2 | 1.4x10<sup>-9</sup> | Up | 10 | 625 |
|  | Ileum | 0 vs 1 Week | IFN response | 3.4 | 5.5x10<sup>-8</sup> | Up | 18 | 167 |
| **IFIT3** | Abomasum | 1 vs 8 Weeks | IFN-induced antiviral protein | 3.1 | 5x10<sup>-20</sup> | Up | 337 | 439 |
| **IFIT2** | Ileum | 0 vs 1 Week | IFN-induced antiviral protein | 2.0 | 7.7x10<sup>-31</sup> | Up | 76 | 275 |
| **MX2** | Abomasum | 0 vs 1 Week | IFN-induced protein | 2.1 | 7x10<sup>-7</sup> | Up | 19 | 86 |
|  | Rumen | 0 vs 1 Week | IFN-induced protein | 5.4 | 8 x10<sup>-7</sup> | Up | 13 | 443 |
|  | Rumen | 1 vs 8 Weeks | IFN-induced protein | -2.8 | 4.3x10<sup>-3</sup> | Down | 443 | 47 |
| **MX1** | Abomasum | 1 vs 8 Weeks | IFN-induced antiviral protein | 2.1 | 5.5x10<sup>-21</sup> | Up | 1048 | 1307 |
|  | Rumen | 0 vs 1 Week | IFN-induced antiviral protein | 3.6 | 2x10<sup>-5</sup> | Up | 220 | 2256 |
| **IDO1** | Ileum | 0 vs 1 Week | Amino acid catabolism | 4.2 | 2x10<sup>-8</sup> | Up | 88 | 1467 |
| **COL1A1** | Rumen | 1 vs 8 Weeks | Collagen - connective tissue | 2.1 | 3.3x10<sup>-5</sup> | Up | 29269 | 92501 |
|  | Ileum | 1 vs 8 Weeks | Collagen - connective tissue | -2.3 | 8.5x10<sup>-15</sup> | Down | 35506 | 6919 |
| **COL1A2** | Rumen | 1 vs 8 Weeks | Collagen - connective tissue | 2.1 | 7.3x10<sup>-5</sup> | Up | 25919 | 80206 |
|  | Ileum | 1 vs 8 Weeks | Collagen - connective tissue | -2.1 | 5.6x10<sup>-14</sup> | Down | 33384 | 7541 |
| **HERC6** | Rumen | 1 vs 8 Weeks | Ubiquitin ligase activity | -3.9 | 4.3x10<sup>-3</sup> | Down | 319 | 16 |
|  | Ileum | 0 vs 1 Week | Ubiquitin ligase activity | 3.2 | 7.7x10<sup>-11</sup> | Up | 25 | 211 |
|  | Abomasum | 1 vs 8 Weeks | Ubiquitin ligase activity | 3.0 | 1.1x10<sup>-11</sup> | Up | 357 | 636 |
| **KRT36** | Rumen | 1 vs 8 Weeks | Keratin | 3.9 | 6.5x10<sup>-8</sup> | Up | 6384 | 69503 |
| **HMGCS2** | Rumen | 0 vs 1 Week | Ruminal ketone body synthesis | 8.4 | 2.3x10<sup>-50</sup> | Up | 66 | 19467 |
| **DUOXA2** | Rumen | 0 vs 1 Week | Thyroid hormone synthesis | 4.1 | 3.2x10<sup>-5</sup> | Up | 12 | 168 |
| **SLC14A1** | Rumen | 0 vs 1 Week | Nitrogen and Iodine Recycling | 2.4 | 9.3x10<sup>-16</sup> | Up | 2181 | 10158 |

Using Venn diagrams (Figure S2) we compared whether any differentially expressed genes were shared across tissues and developmental stages. Relatively few genes were shared between developmental stages (0-1 week and 1-8 weeks), fifty genes were shared between the rumen time points, 19 between the ileum and 13 between the abomasum (Figure S2). A greater number of differentially expressed genes were shared between the rumen and ileum than either of these two tissues and the abomasum, which reflects the different functional roles of these three tissues. Relatively few genes were differentially expressed in two or more tissues and time points, the most notable being *HERC6* which is involved in ubiquitin ligase activity and *ISG15* an IFN-gamma-inducing cytokine playing an essential role in antimycobacterial immunity (Table 3).

The structural composition of the rumen and ileum changes significantly during the transition from pre-ruminant to ruminant, with the most dramatic physical changes associated with the rumen epithelium (Baldwin *et al.* 2004). Several genes associated with connective tissue and collagen were differentially expressed between one week and 8 weeks of age. Genes exhibiting greater than 2-fold up-regulation included *COL1A1* and *COL1A2* (Table 3). Genes associated with keratinocytes and the epithelial signature of the rumen exhibited differential expression patterns according to developmental stage. *KRT36*, for example, showed a more than 4-fold increase in expression between one and 8 weeks of age (Table 3). *KRT36* has been shown to exhibit significant transcriptional responses to changes in diet in dairy cattle (Li *et al.* 2015) and the change in diet between one and 8 weeks of age is likely to be driving at least a proportion of the observed expression patterns.

Similarly, the mitochondrial enzyme encoding gene *HMGCS2*, which belongs to the HMG-Co synthase family and catalyses the first reaction in ketogenesis (Lane *et al.* 2002) was 8-fold up-regulated in the rumen between birth and one week of age (Table 3). *HMGCS2* has also been shown to be differentially expressed in the calf rumen during the transition from pre-ruminant to ruminant (Connor *et al.* 2013; Kato *et al.* 2016) and in the rumen of developing lambs (Lane *et al.* 2002; Wang *et al.* 2016). It has been suggested that dietary changes through early development are likely to promote ketogenesis in rumen epithelial cells via PPAR-α-mediated activation of *HMGCS2* to promote papillary development, as well as activation of genes promoting fatty acid beta-oxidation to support cellular differentiation (Connor *et al.* 2013). *SLC14A1* has also been shown to be differentially expressed in the calf rumen during the transition from pre-ruminant to ruminant (Connor *et al.* 2013) and was differentially expressed in the rumen of lambs between birth and one week of age (Wang *et al.* 2016). It encodes the protein SLC14A1 which mediates the basolateral cell membrane transport of urea, a key process in nitrogen secretion into the ruminant GI tract (Abdoun *et al.* 2009). A characteristic of ruminants is a high level of nitrogen recycling in the GI tract. Recycling of nitrogen via urea secretion into the rumen allows ruminants to survive on low-protein diets whilst at the same time producing milk and meat for human consumption (Abdoun *et al.* 2009).

### Large differences in developmental stage specific expression patterns are associated with the immune response

Several genes involved in the immune response were differentially expressed in the rumen, ileum and abomasum of developing lambs during the transition from pre-ruminant to ruminant. Many of these genes are likely to be part of the acute phase immune response, by regulating production of key cytokines such as *IL-6* and thus mediating activation of the NF-κB signaling pathways. Nuclear factor (NF)-κB and inhibitor of NF-κB kinase (IKK) proteins regulate innate- and adaptive-immune responses and inflammation (Perkins 2007). *IL36A* and *IL36B* are thought to influence the skin inflammatory response by acting directly on keratinocytes and macrophages and indirectly on T-lymphocytes to drive tissue infiltration, cell maturation and cell proliferation (Foster *et al.* 2014). *IL36A* is a cytokine that can activate the NF-kappa-B and MAPK signaling pathways to generate an inflammatory response and shows almost a 4-fold increase in expression between one and 8 weeks of age in the rumen. Other C-type lectins and genes involved in cytokine signalling also showed differential expression between the different developmental stages. *CLEC4E*, for example, was more than 2-fold upregulated in the Ileum between one and 8 weeks of age. *JAK3*, which encodes a member of the Janus kinase (JAK) family of tyrosine kinases is involved in cytokine receptor-mediated intracellular signal transduction (Yeh & Pellegrini 1999), and was more than 5-fold upregulated in the rumen between birth and one week of age.

Other immune genes, including *PIP* (Prolactin Induced Protein), were upregulated 3-fold in the rumen of lambs between one and 8 weeks of age. *PIP* has been shown to be preferentially expressed in the rumen of adult sheep (Xiang *et al.* 2016a), and is thought to play a role in mucosal immunity in ruminants (Hassan *et al.* 2008). Multiple IFN-inducible genes, including *IFIT2*, *IFIT3*, *MX1*, *MX2* and *ISG15*, were differentially expressed in the abomasum, rumen and ileum during the transition from pre-ruminant to ruminant (Table 3). Three of these genes - *IFIT2*, *IFIT3* and *MX1* - have recently been shown to be differentially expressed in sheep fibroblast cells in a type I IFN-induced antiviral state (Shaw *et al.* 2017). The expression pattern of some of these genes varied through development. *MX2* for example, is up-regulated in the rumen between birth and one-week of age and down-regulated between one and eight weeks of age (Table 3). The differential expression patterns of these genes throughout the development of the rumen, abomasum and ileum highlights both their importance in the innate immune response and the role of the GI tract as an immunologically active site.

Several genes involved in both metabolism and immunity were differentially expressed during the transition from pre-ruminant to ruminant. *DUOXA2*, for example is involved in thyroid hormone synthesis and lactoperoxidase-mediated antimicrobial defense at the surface of mucosa (Bae *et al.* 2010). The rumen is the main site of colonisation by micro-organisms as the lamb develops. *DUOX2* and *DUOXA2*, which encode subunits of dual oxidase have previously been shown to be upregulated in the rumen of adult sheep (Xiang *et al.* 2016a; Xiang *et al.* 2016b) and here we found *e*xpression of *DUOXA2* 4-fold up-regulated in the rumen between birth and one week of age. *DUOXA2* might be involved in controlling microbial colonization as the lamb transitions from pre-ruminant to ruminant. Similarly, *IDO1* was more than 4-fold upregulated in the ileum of lambs between birth and one week of age and has been implicated in immune modulation through its ability to limit T-cell function and engage mechanisms of immune tolerance (Grohmann *et al.* 2003; Plain *et al.* 2011). *IDO1* encodes indoleamine 2,3-dioxygenase (IDO) which is a heme enzyme that catalyzes the first and rate-limiting step in tryptophan catabolism to N-formyl-kynurenine (Grohmann *et al.* 2003). Through its expression in monocytes and macrophages this enzyme modulates T-cell behaviour by its peri-cellular catabolization of the essential amino acid tryptophan (Munn & Mellor 2013). It has also been shown to be highly expressed in the jejunal mucosa of pre-weaning calves (Hammon *et al.* 2018).

### Comparative analysis of the rumen, colon and ileum of one week old age-matched sheep and goats reveals differences in expression of genes involved in metabolism and immunity

We performed a comparative analysis of the gene expression estimates for age-matched one week old sheep and goats for three GI tract tissues: rumen, ileum and colon. The goat gene expression estimates as transcripts per million (TPM) both for these tissues and alveolar macrophages are included in Table S9. Full lists of all differentially expressed genes between sheep and goat are included in Table S10 (rumen), Table S11 (ileum) and Table S12 (colon). The top 25 differentially expressed genes between sheep and goat in either direction for each of the three tissues are shown in Figure 7. Differentially expressed genes between the rumen of sheep and goats included several solute carrier genes (*SLC5A8*, *SLC9A4, SLC14A1*, and *SLC27A6*), a myosin V heavy chain gene *MYO5A* associated with connective tissue, and *KRT3* which is involved in the simple and stratified differentiation of epithelial tissues. Other genes within these super-families were also differentially expressed between the ileum and colon of sheep and goats. Similarly, genes associated with immunity and metabolism were differentially expressed between sheep and goat GI tract tissues. For example, C-C motif chemokine ligands *CCL5* and *CCL20* were upregulated, respectively, in the goat ileum and colon. Chemokines form a superfamily of genes involved in immunoregulatory and inflammatory processes (Griffith *et al.* 2014). *CCL5* is one of the predominant cytokines expressed during damage and inflammation of epithelial keratinocytes (Wetzler *et al.* 2000). In contrast, *CSF3R* was down regulated in the colon of goats relative to sheep (Figure 7). *CSF3R* (also known as *GCSF*) encodes a protein that controls the production, differentiation and production of granulocytes (Nagata & Fukunaga 1991). Differences in expression between the two species might therefore reflect variation in the density of granulocyte cells. *CSF3R* has been shown to undergo changes in anatomical distribution during human fetal development in the intestine (Calhoun *et al.* 1999), and similar trends could also occur in neonatal animals.

**Figure 7:**
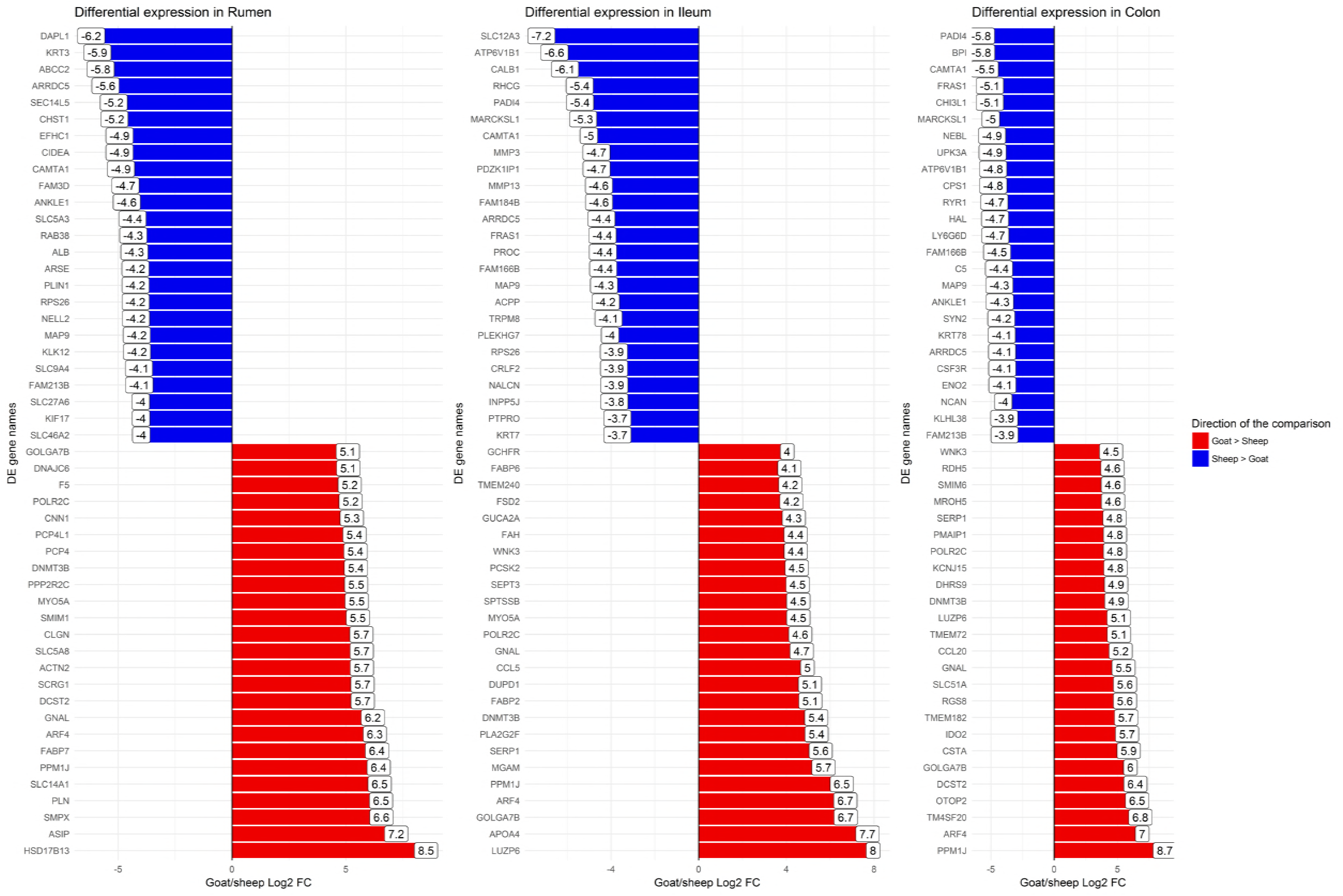
The top 25 differentially expressed genes up-regulated and top 25 down-regulated genes between goat and sheep in the rumen, ileum and colon of age-matched one-week old animals.

Other genes with key roles in metabolic pathways and immune regulation were differentially expressed between goats and sheep, including *IDO2*, which was upregulated in the colon of one-week old goats relative to age-matched sheep. *IDO2*, with its paralogue *IDO1,* mentioned above, catalyses the first and rate-limiting step of the catabolism of the essential amino acid tryptophan along the kynurenine pathway (Grohmann *et al.* 2003) and is involved in the metabolic control of immune responses (Munn & Mellor 2013). IDO has been implicated in downregulating immune responses to *Mycobacterium avium* subsp. *paratuberculosis*, the causative agent of Johne’s disease (Plain *et al.* 2011). In sheep and cattle an increase in IDO expression correlates with progression to clinical mycobacterial disease (Plain *et al.* 2011) with the small intestine the primary site of infection (Koets *et al.* 2015). Since the GI tract contains important regions related to immunity differences in the expression of genes with immune and metabolic function in the tissues analysed might underlie differences in disease susceptibility and response to pathogens between the two species. Interestingly, *SLC14A1* which was differentially expressed in the rumen of sheep between birth and one week of age, was also up-regulated in the rumen of goats relative to sheep (Figure 7). As mentioned above the protein encoded by *SLC14A1* mediates the basolateral cell membrane transport of urea, a key process in nitrogen secretion into the ruminant GI tract (Abdoun *et al.* 2009). Nitrogen recycling is an important topic in ruminant management because movements of nitrogen across the GI tract can more than double nitrogen intake and have major effects on nitrogen metabolism (Lapierre & Lobley 2001). Nitrogenous content of urea between sheep and goats has been shown to vary (Bristow Andrew *et al.* 1992), which may be influenced by species specific differences in expression of key genes involved in iodine and nitrogen recycling pathways in ruminants.

The results from our differential expression analysis of the three GI tract tissues in age-matched sheep and goats indicate that there are basal differences in the expression of some key genes involved in pathways related to immunity and metabolism between the two species. These genes would be candidates for further analysis to elucidate the functional significance of species specific differences in gene expression e.g. (Young *et al.* 2018).

## Conclusions

By characterising tissue specific transcription in the GI tract through the transition from pre-ruminant to ruminant we have shown that there are significant developmental stage specific differences in gene expression particularly between neonatal lambs and 8 week and adult sheep. These differences were most obvious in the rumen complex where significant morphological and physiological changes occur as the lamb transitions from a milk-based to a grass and pellet diet. Differences in the expression of protein coding genes with age were observed both when the whole transcriptome was included in the analysis, and also when only a subset of macrophage specific genes were analysed.

We focused only on protein-coding genes in this study, and have presented a wider characterisation of non-coding transcripts elsewhere (Bush *et al.* 2018). A low level, highly tissue specific expression pattern is characteristic of lncRNAs in sheep and goats (Bush *et al.* 2018). Further characterisation of the lncRNAs and other non-coding transcripts expressed specifically in the GI tract through development, might help to infer something about their function and would be an interesting direction for future work. Similarly, analysis of isoform regulation using RNA-Seq data, as in (Katz *et al.* 2010), in the GI tract tissues would also be interesting, although both this and analysis of the non-coding transcriptome would require further sequencing to generate high depth total RNA-Seq libraries.

Other studies have examined links between feeding regime, the host transcriptome and bacterial diversity in sheep using sequencing analysis of 16S rRNA genes (Wang *et al.* 2016). A full characterisation linking tissue- and developmental stage-specific microbial colonisation of the GI tract would require a detailed 16S sequencing metagenomic approach e.g. (Wallace *et al.* 2015). Further work would ideally examine this link further to profile in parallel transcription in the GI tract and microbial colonisation to provide information as to how microbial colonisation influences transcription during development. An additional area to explore further would be the relative *IL10* insufficiency in the GI tract of the lambs – which suppresses the macrophage response to intestinal microbes – as this could potentially be beneficial to ruminant species which, compared to monogastric mammals, have a quite different relationship with gut microbes.

The results we present in this study lay a foundation for further work, providing baseline estimates of gene expression in the GI tract at the whole tissue level from healthy lambs. In their 2002 review of gene expression in the ruminant GI tract Connor *et al.* indicated that the use of techniques such as laser-capture microdissection will be needed to further characterize expression profiles of individual cell types within the GI tract, and to remove expression biases that may occur in studies evaluating whole tissue samples (Connor *et al.* 2010). Cutting edge single cell RNA-Seq technology provides the solution to this, allowing a cell specific level of resolution of gene expression profiles. Single cell messenger RNA-Seq has already been applied to cells from mouse GI tract organoids revealing rare cell types (Grün *et al.* 2015) and the technology will hopefully be applied in the future to cells from the GI tract of ruminants, since *in vitro* systems are now available (Hamilton *et al.* 2018).

One of the most significant physiological challenges to neonatal and juvenile ruminants is the development and establishment of the rumen. The transition from preruminant to ruminant involves not only growth of the rumen and cellular differentiation but also has a major effect on metabolism, immunity and physiological processes in other GI tract tissues (Baldwin *et al.* 2004), which is reflected in the extensive transcriptional complexity observed in this study. Using this sub-set of RNA-Seq data, from our high resolution of atlas of gene expression in sheep (Clark *et al.* 2017b) we have improved our understanding of the genetic and genomic mechanisms involved in the transition between pre-ruminant and ruminant in sheep and highlighted key genes underlying healthy growth and development that could be utilised to improve productivity in sheep and other ruminants.

## Author’s Contributions

ELC coordinated and designed the study with assistance from MEBM, ALA and DAH. DAH acquired the funding with ALA. MEBM, ILF and ELC performed sample collection from sheep. CM, ELC and ZML performed sample collection from goats. GMD and CM performed the RNA extractions. ELC performed the network cluster and gene expression analysis. MS performed the Principal Component Analysis and created Figure 1. CM performed the comparative analysis of sheep and goat. DAH and ZML provided guidance on interpretation of the gut macrophage transcriptional signatures. SJB performed all bioinformatic analyses. ELC wrote the manuscript. All authors read and approved the final manuscript.

## Acknowledgements

The authors would like to thank the farm staff at Dryden farm and members of the sheep tissue collection team from The Roslin Institute and R(D)SVS who were involved in tissue collections for the sheep and goat gene expression atlas projects. The authors are also grateful for the support of the FAANG Data Coordination Centre in the upload and archiving of the sample data and metadata and BioGPS for hosting the sheep atlas dataset on their annotation portal.

## Supplemental Material

**Table S1:** Averaged TPM expression estimates for sheep GI tract tissues, alveolar macrophages, thoracic oesophagus and liver, including the TxBF and Texel data.

**Table S2:** List of genes within each cluster from the gene-to-gene network graph (Figure 2) including sheep GI tract tissues, alveolar macrophages, thoracic oesophagus and liver.

**Table S3:** GO term enrichment for molecular function, cellular component and biological process for each cluster from the gene-to-gene network graph for sheep (Figure 2).

**Table S4:** Individual sheep GI tract TPM expression estimates used for this analysis, which is a subset of data from the TxBF sheep atlas.

**Table S5:** Differentially expressed genes in the sheep rumen using pairwise comparisons between birth and one week and one week and 8 weeks of age.

**Table S6:** Differentially expressed genes in the sheep ileum using pairwise comparisons between birth and one week and one week and 8 weeks of age.

**Table S7:** Differentially expressed genes in the sheep abomasum using pairwise comparisons between birth and one week and one week and 8 weeks of age.

**Table S8:** GO term enrichment of differentially expressed genes in the rumen, ileum and abomasum of sheep between two developmental stages birth and one week and one week and 8 weeks of age.

**Table S9:** Individual TPM expression estimates for rumen, ileum and colon from one-week old goats.

**Table S10:** Differentially expressed genes in rumen samples from one-week old sheep and goats.

**Table S11:** Differentially expressed genes in ileum samples from one-week old sheep and goats.

**Table S12:** Differentially expressed genes in colon samples from one-week old sheep and goats.

**Figure S1: Principal Component Analysis of sheep GI tract samples illustrating separation by tissue (A & B) and developmental stage (C & D) in three components.**

**Figure S2: Visualisation of differentially expressed genes within and between ileum, rumen and abomasum tissues at two developmental stages (0-1 week) and (1-8 weeks) in sheep using Venn diagrams.**

